# Pushing Sensitivity and Specificity Limits in Native Structural Biology: ^19^F Multinuclear Dynamic Nuclear Polarization with Magic Angle Spinning

**DOI:** 10.1101/2025.07.16.664958

**Authors:** Kumar Tekwani Movellan, Daniel Banks, Christian Reiter, James Kempf, Angela M. Gronenborn, Tatyana Polenova

## Abstract

Understanding protein structures and their interactions within natural cellular environments is essential for deciphering cellular processes and advancing therapeutic development. Obtaining atomic-level information about protein structural changes in cellular contexts poses a significant challenge. Here, we introduce a ^19^F-based, ^1^H-assisted dynamic nuclear polarization (DNP) magic angle spinning (MAS) NMR approach that offers exceptionally high sensitivity and specificity, enabling background-free detection of target proteins in mammalian cells for atomic-level structural analysis. We demonstrate this methodology in A2780 cells for the human Cyclophilin A (CypA) protein with a single fluorine atom incorporated in the sole tryptophan residue. We achieved significant sensitivity gains through ^1^H-^19^F cross-polarization (CP), with subsequent ^19^F-^13^C double CP providing unique structural information. Remarkably, using ^1^H-^19^F^13^C magnetization transfer allowed selective detection of ^13^C signals from CypA residues up to 6 Å away from the fluorine label. Taken together, our study establishes a framework for investigating protein structure, dynamics, and interactions in mammalian cells by DNP MAS NMR.

## INTRODUCTION

Protein-protein interactions are crucial to cell-cell communication and chemical reactions within the cell. Understanding these interactions in cellular contexts is essential for elucidating biological mechanisms in both healthy and disease states, as well as for developing effective therapeutic strategies. However, gaining atomic-level insights into protein structure, dynamics, and interactions in cellular environments remains challenging with current methodologies due to their limited sensitivity and/or specificity.^1^ Multiple research teams have explored solution nuclear magnetic resonance (NMR) spectroscopy^2–5^ or dynamic nuclear polarization (DNP) magic angle spinning (MAS) solid-state NMR to address these challenges.^6–8^ NMR spectroscopy is unparalleled in providing atomic-level information on both the structure and dynamics of biomolecules, with exquisite specificity across a wide range of time scales. Concurrently, the sensitivity gains offered by DNP create opportunities for detecting proteins at their endogenous concentrations within cells.^9^ Since the pioneering studies by Serber et al.^2^ and Selenko et al.^3^ in the 2000s, in-cell solution NMR has gained increasing popularity.^10^ Simultaneously, several research groups have explored DNP magic angle spinning (MAS) solid-state NMR for in-cell applications.^6,7,11–13^ While powerful, most in-cell NMR studies reported to date rely on ^13^C and ^15^N-based experiments involving isotopically enriched proteins, and the resulting spectra are often plagued by natural abundance cellular background signals and/or signal loss due to protein interactions with cellular components.^5,6^

A powerful alternative to traditional 2D ^13^Cand ^15^N correlation spectra is using ^19^F as a site-specific probe for in-cell solution and MAS NMR applications,^5,14,15,7,16,17^ exploiting its favorable NMR properties.^18,19^ First, fluorine can be easily incorporated into proteins biosynthetically using fluorinated amino acids. Second, the fluorine’s virtual absence in biological systems renders ^19^F NMR spectra background-free. And third, the large chemical shift range of ^19^F, which exceeds 300 ppm combined with the sitespecific incorporation, reduces problems associated with resonance overlap. Recently, we reported a ^19^F-based fast MAS DNP NMR approach for in-cell applications.^7^ While DNP signal enhancements ranging from 30 to 40-fold could be achieved for ^19^F resonances through direct polarization mechanism, these experiments were hampered by long signal buildup times exceeding 10 s. Furthermore, the polarization agent had to be electroporated into cells together with the protein, resulting in its rapid reduction.

Here, we present an alternative ^19^F in-cell DNP MAS NMR strategy based on double and triple resonance ^1^H-^19^F and (^1^H-^19^F)-^13^C experiments conducted at a moderate MAS frequency of 12 kHz. This approach employs an indirect DNP transfer pathway via ^1^H-^19^F cross-polarization (CP) with modest concentrations of AMUPol biradical added extracellularly. Since AMUPol does not cross the cellular membrane, its rapid reduction inside cells does not occur, and ^1^H-^1^H spin diffusion can be taken advantage of to propagate the polarization from outside to inside the cells. As a result, even greater signal enhancements than those observed in our previous ^19^F direct-DNP experiments were obtained. Importantly, these additional sensitivity gains facilitate ^19^F-^13^C magnetization transfers that yield critical structural information in spectra completely devoid of cellular background signals. We present results on human Cyclophilin A (CypA), a 18 kDa protein that contains a single tryptophan residue (Fig. 1), delivered into mammalian cells by electroporation. We estimated >50-fold increases in ^19^F resonance intensities in the incell ^1^H-assisted ^1^H-^19^F CP DNP-enhanced MAS NMR experiments, enabling the detection of low-nanomole quantities of 4F-Trp-U-^13^C,^15^N-CypA with a signal-to-noise ratio (SNR) of 9.6 in spectra acquired in approximately 1.7 hours. Furthermore, using (^1^H-^19^F)-^13^C dipolar-based experiments applying two consecutive CP transfer steps allowed us to eliminate the natural-abundance ^13^C cellular background and detect signals arising solely from CypA carbon atoms within 6 Å of the fluorine atom. Our results also highlight the importance of a dedicated ^1^H channel in HFX DNP probes for polarization transfers and decoupling. More broadly, our approach paves the way for probing protein conformational changes and interactions around judiciously placed fluorine atoms in mammalian cells.

**FIGURE 1.**
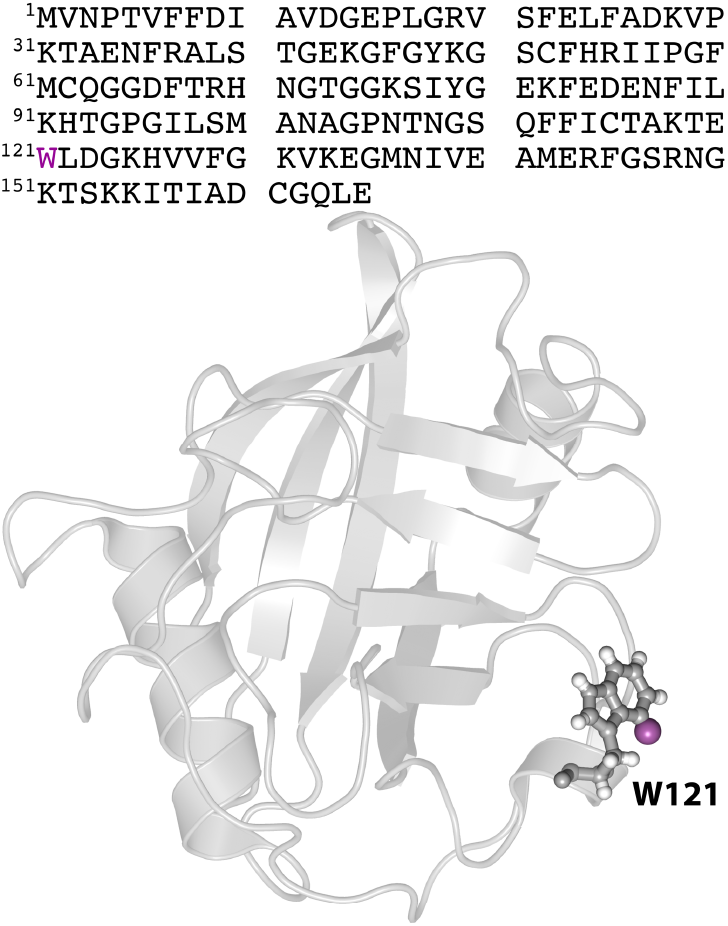
Amino acid sequence and structure of CypA. Top: The amino acid sequence of CypA with the W121 residue labeled in magenta. Bottom left: Ribbon representation of the CypA structure (PDB: 3K0M). The side chain of W121 is depicted in stick representation and the fluorine atom at position 4 of the indole ring is shown as a magenta sphere.

## RESULTS AND DISCUSSION

### Sensitivity and spectral resolution of *in-cell* ^19^F ^1^H-assisted DNP-enhanced MAS NMR experiments

We delivered 4F-Trp,U-^13^C,^15^N-CypA into human A2780 cells via electroporation, while AMUPol was added extracellularly after cell recovery following electroporation. This protocol is advantageous since AMUPol remains outside the cell and is not reduced rapidly, allowing for ^1^H-^1^H spin diffusion to propagate the polarization from outside to inside the cells, via a relayed DNP mechanism. The sample contained 20 million cells, with 554 μM (12.2 nanomoles) of electroporated CypA protein (Fig. S1, Supporting Information) and 6.4 mM (31 micromoles) of AMUPol in the MAS rotor (Table 1). The ^1^H-^19^F DNP-enhanced CPMAS NMR spectrum acquired at a MAS frequency of 12 kHz is presented in Fig. 2A (top trace). Signalto-noise ratios (SNR) of 9.6 (determined with respect to the isotropic peak intensity, SNR^I^) and 1.4*10^5^ (determined using the integrated intensities summed over the isotropic and spinning sideband resonances, SNR^int^) were reached after 1.7 hours of signal averaging (Table S1, Supporting Information). The integrated normalized SNR value, SNR^int^_norm_, defined as the SNR^int^ per square root of experiment time, per nanomole of protein, per number of fluorine atoms in the protein, was 1134. In contrast, very weak signals were detected in the control “microwave-off” experiment after 6.8 hours of signal averaging. Given that the SNR from the isotropic peak did not exceed 1.4 in the control experiment (Fig. S2, Supporting Information), we could not accurately measure the ^19^F DNP signal enhancements. Therefore, we estimated the enhancements as a ratio of SNR^int^ for the mw-on and mw-off spectra, with the signal and noise regions taken to be the same as in the DNP-enhanced spectrum (Fig. S3, Supporting Information). According to this protocol, we estimated the DNP enhancements, ε, to exceed 50-fold. These values most likely are in agreement with the ε value of approximately 40, accurately determined from ^1^H-^13^C CPMAS ^13^C-detected DNP MAS spectra (Fig. S4, Supporting Information).

**Table 1.** Summary of sample conditions and signal-to-noise ratio (SNR) in ^1^H-assisted ^19^F DNPenhanced MAS NMR experiments on 4F-Trp,U-^13^C,^15^N-CypA in human A2780 cells.

|  |  |
| --- | --- |
| Cells/rotor ($\times 10^6$) | 20 |
| Protein concentration in the NMR sample ($\mu\text{M}$) | 554 |
| Amount of protein in the NMR sample (nanomoles) | 12.2 |
| Active rotor volume ($\mu\text{L}$) | 30 |
| [AMUPol] by EPR (mM) | 6.4 |
| Buffer | 10% DMSO-$\text{d}_6$, 20% $\text{D}_2\text{O}$, 20% FBS, 50% RPMI |
| Cell viability (%) | >90 |
| $^1\text{H}$ $T_b$ (s) | 1.5/25 |
| $^{\text{a,d}}\text{SNR}^{\text{I}}$ | 9.6 |
| $^{\text{b,d}}\text{SNR}^{\text{int}}$ | $1.4 \times 10^5$ |
| $^{\text{c,d}}\text{SNR}_{\text{norm}}^{\text{int}}$ | 1134 |
| Experiment time (min) | 102 |

$^{\text{a}}\text{SNR}^{\text{I}}$ is calculated as a ratio of isotropic peak intensity to noise RMSD.
$^{\text{b}}\text{SNR}^{\text{int}}$ is calculated as a ratio of the integrated signal intensity over all spinning sidebands to noise RMSD.
$^{\text{c}}\text{SNR}_{\text{norm}}^{\text{int}}$ is calculated as a ratio of the integrated signal intensity over all spinning sidebands to noise RMSD, per number of nanomoles per the square root of the experimental time.
The $\text{SNR}^{\text{I}}$ , $\text{SNR}^{\text{int}}$ , and $\text{SNR}_{\text{norm}}^{\text{int}}$ were determined from the $^1\text{H}$ - $^{19}\text{F}$ DNP CPMAS spectrum acquired for 4.9 ms and processed without apodization.

**FIGURE 2.**
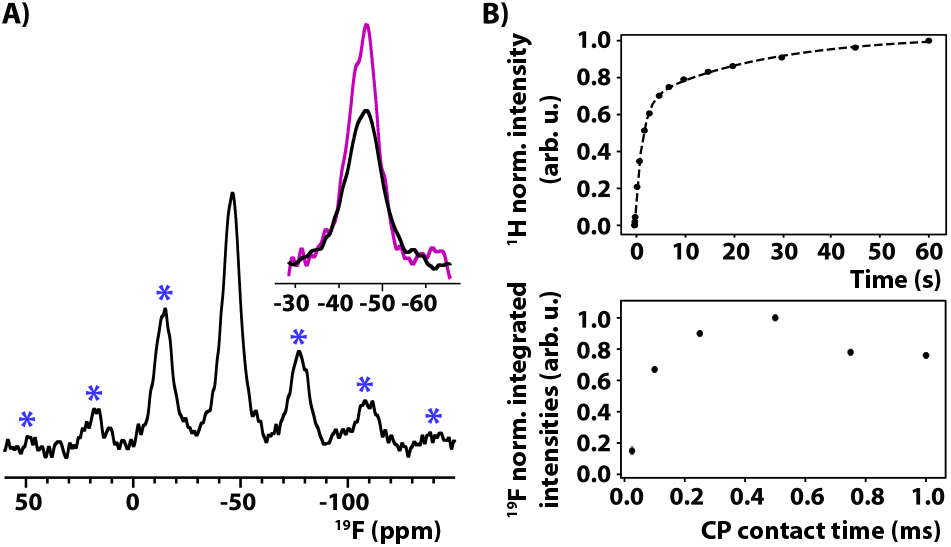
Sensitivity of in-cell DNP-enhanced ^1^H-^19^F CPMAS NMR experiments of 4F-Trp,U^13^C,^15^N-CypA. **(A)** In-cell DNP-enhanced spectrum of a 4F-Trp,U-^13^C,^15^N-CypA sample containing ∼20 million cells, 554 μM CypA protein and 6.4 mM AMUPol. The region around the isotropic peak is expanded in the inset for the spectra recorded with (magenta trace) and without (black trace) ^1^H decoupling. Spectra were acquired in 103 min with 2048 scans and a recycle delay of 3 s. The spinning sidebands are marked with blue asterisks. **(B)** ^1^H DNP signal buildup (top) and ^1^H-^19^F CPMAS signal intensity vs. contact time (bottom). Other acquisition and processing parameters are provided in Table S2, Supporting Information. The active rotor volume was 30 μL.

The ^1^H DNP signal buildup clearly exhibits bi-exponential behavior, with corresponding buildup times (T_b_) of 1.5 and 25 s (Fig. 2B) with contributions of ∼62 and 38 % respectively, which align with previously reported values from in-cell DNP studies.^13^ Since DNP buildup times are determined by the electron-^1^H polarization transfer rates, this suggests the presence of two pools of protons: one close to and another further away from the biradical. As AMUPol was added extracellularly following electroporation and after cell recovery, no sizable concentration of biradical should exist inside the cells, and the fast and slow T_b_ likely represent protons outside and inside the cells, respectively.

In contrast, as we found in our previous study, the in-cell ^19^F DNP signal buildup in direct-polarization experiments is on the order of 13-16 s.^7^ In that study, the probe could only be operated in a single-resonance mode, either ^19^F or ^1^H. To perform the experiments in a reasonable amount of time, we opted for a compromise and used a recycle delay of 5 s. With the current HFX DNP probe configuration, we could take advantage of the fast ^1^H DNP signal buildup.

Fig. 2B shows the signal intensity as a function of the ^1^H-^19^F CP contact time. The maximum signal is observed at 0.5 ms, at a significantly shorter contact time than seen in ^1^H-^19^F CPMAS experiments performed at temperatures above 0 ºC without DNP enhancement.^20–22^

The ^19^F line widths of the isotropic peak are approximately 9 and 7 ppm in the absence and presence of ^1^H decoupling, respectively (Fig. 2A inset). Additionally, the ^1^H-decoupled spectrum exhibits a gain in signal intensity of approximately 20-25%. Although the linewidth with decoupling is smaller than without, a 7 ppm wide line is surprisingly large, given that the signal originates from a single 4F-W121 site. In addition, the linewidths are also significantly larger than the homogeneous line widths of 2-3 ppm previously observed in ^19^F DNP MAS experiments on CA assemblies *in vitro*^23^ and N^NTD^ in cells.^7^ We hypothesize that the ^19^F resonance of 4F-Trp,U-^13^C,^15^N-CypA is inhomogeneously broadened due to interactions with other cellular binding partners, as suggested by a previous ^19^F in-cell solution NMR study.^5^ Recently, we reported 30to 40-fold ^19^F signal enhancements in direct-polarization DNP experiments performed at a MAS frequency of 40 kHz on SARS-CoV-2 N^NTD^ protein in A2780 mammalian cells.^7^ In this case, the sample contained approximately 1.2 to 1.5 million cells with 200 μM of N^NTD^ and 13 mM AMUPol. Here, in contrast, no direct-polarization ^19^F signal was observed after 4.5 hours of signal averaging in the current DNP experiments on 4F-Trp,U-^13^C,^15^N-CypA carried out at a MAS frequency of 12 kHz (Fig. S2 of the Supporting information). This result is not surprising, given that the current sample consists of only 6.4 mM AMUPol delivered extracellularly, highlighting the critical role of ^1^H-assisted DNP mechanism for achieving sensitivity gains in ^1^H-^19^F cross-polarization DNP-based experiments at moderate spinning frequencies. The rapid buildup of the ^1^H DNP signal, which occurs within 1.5 seconds, allows for recycle delays of 2 to 3 seconds in ^1^H-^19^F CPMAS DNP experiments, compared to the 15 seconds required in ^19^F direct-DNP experiments. These sensitivity gains, along with the capability for ^1^H decoupling and/or heteronuclear polarization transfers, are only possible if a dedicated ^1^H channel is available, as in the current HFX DNP probe, as discussed below.

### ^19^F chemical shift anisotropy

Fluorosubstituted tryptophans display significant ^19^F chemical shift anisotropies (CSAs). The experimentally determined reduced anisotropy parameters, δ_σ_, in crystalline and powder samples range from 47.4 to 67.6 ppm, depending on the fluorine position in the indole ring, the crystal form, and the hydration level.^24^ The maximum achievable MAS frequency in this study is 12 kHz, limited by the 3.2 mm HFX probe design. This spinning frequency is insufficient for fully averaging out the CSA interaction. Indeed, spinning sidebands (SSBs) are prominent in the spectra of 4F-Trp,U-^13^C,^15^N-CypA, as illustrated in Fig. 2A. Although the incomplete ^19^F CSA averaging is not ideal with regard to sensitivity, since only 48% of the integrated signal intensity resides in the isotropic peak, the moderate spinning frequency permits the determination of the ^19^F CSA tensor parameters by directly fitting the SSBs (Table S1, Supporting Information). The resulting δ_σ_ value is 67±4 ppm, and the asymmetry parameter, η, is 0.5±0.4. These values are generally consistent with those reported for crystalline 4F-Trp.^24^

### ^19^F-filtered (^1^H-^19^F)-^13^C CPMAS spectra: local structure

While the large ^19^F signal enhancements achieved in direct-DNP^7^ and ^1^H-^19^F CPMAS-DNP experiments (this study) offer a powerful tool for studying proteins in cells, gaining detailed atomic-level conformational information requires detecting signals from other atoms in the protein, such as ^13^C. Unfortunately, ^13^C-detected in-cell NMR spectra, whether in solution or solids, without or with DNP enhancements, contain signals arising from all carbon-containing species in the sample and are typically dominated by the natural-abundance carbon signals.^6^ Indeed, as shown in Fig. 3A (bottom trace), the DNP-enhanced ^1^H-^13^C CPMAS spectrum for the in-cell 4F-Trp,U-^13^C,^15^N-CypA sample under investigation is overwhelmed by the natural abundance ^13^C signals and closely resembles spectra reported previously for other in-cell samples.^6,12,25^ To eliminate the natural-abundance ^13^C background signals and selectively reveal peaks arising from 4F-Trp,U-^13^C,^15^N-CypA, we used ^19^F as a filter in a (^1^H-^19^F)-^13^C DNP-enhanced CPMAS experiment. The corresponding spectrum is shown in Fig. 3A (top trace) and contains only 4F-Trp,U-^13^C,^15^N-CypA resonances. Notably, the ^13^C peak widths are 2-4 ppm. Moreover, since CypA’s 3D structure is intact in the cells,^5^ its ^13^C chemical shifts are known (BioMagResBank, BMR27265, Tables S6-S10). Given that 4F-Trp,U-^13^C,^15^N-CypA contains only a single fluorine atom, we were able to tentatively assign the peaks in the spectrum by inspection. Gratifyingly, the ^13^C chemical shifts correspond to those for the backbone and side chain carbon atoms belonging to three residues, T119-W121, confined to a sphere of 6 Å around the fluorine atom (Fig. 3A, B). We note that, while the carbon atoms of the 4FW121 indole ring are also within 6 Å of the fluorine atom, they are natural abundance carbons because of our labeling scheme and, therefore, were not detected in the spectra. Although the sensitivity of this (^1^H-^19^F)-^13^C CPMAS spectrum is low, taking 51 hours for its acquisition, such kinds of experiments pave the way for in-cell ^19^F DNP-enhanced heteronuclear correlation spectroscopy.

**FIGURE 3.**
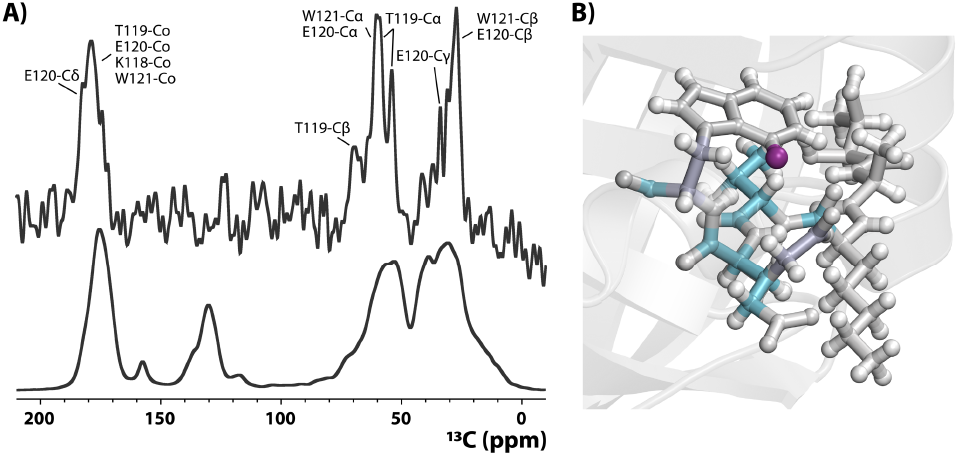
Probing the local protein environment by ^19^F-filtered in-cell ^13^C-detected DNP-enhanced spectroscopy. **(A)** In-cell DNP-enhanced (^1^H-^19^F)-^13^C CPMAS spectrum (top) ^1^H-^13^C CPMAS spectrum (bottom) of a 4F-Trp,U-^13^C,^15^N-CypA sample containing ∼20 million cells, 554 μM CypA, and 6.4 mM AMUPol. The (^1^H-^19^F)-^13^C spectrum was acquired with 61,440 scans for a total of 51 hrs. The ^1^H-^13^C spectrum was acquired with 256 scans for a total of 0.2 hrs. For both spectra, the recycle delay was 3 s, and the MAS frequency was 12 kHz. **(B)** Local structure near the W121 residue of CypA (PDB: 3K0M). The W121 side chain is shown in stick representation, and the fluorine atom is depicted by a magenta sphere. The carbon atoms of K118, T119, E120, and W121, for which signals were observed in the (^1^H-^19^F)-^13^C spectrum, are shown as light purple and light cyan spheres corresponding to distances to the fluorine of <4 Å and <6 Å, respectively.

## CONCLUSIONS AND OUTLOOK

The current study establishes the proof of concept for ^19^F-based ^1^H-assisted multinuclear correlation DNP-enhanced MAS NMR spectroscopy of proteins in native cellular environments. The exceptional specificity provided by fluorine, combined with significantly enhanced sensitivity from ^1^H-assisted DNP and decoupling, enables the recording of multinuclear correlation experiments that were previously unattainable without a dedicated ^1^H channel. Notably, substantial DNP signal enhancements, exceeding 28-fold, were achieved with modest, 6.4 mM extracellular AMUPol concentrations. Providing AMUPol outside the cell overcomes problems with rapid biradical reduction in cellular environments.

While the results presented here represent a significant breakthrough, the currently achievable MAS frequency range of up to 12 kHz in the 3.2 mm HFX DNP probe still poses challenges with respect to sensitivity and resolution. To extend DNP-based native structural biology to a broader research community beyond specialists, it is crucial to develop dedicated HFX DNP probes capable of spinning frequencies of at least 40 kHz. Furthermore, to enhance sensitivity and resolution in the ^19^F-based ^1^H-assisted multinuclear DNP experiments, high and ultrahigh magnetic fields, ranging from 14.1 to 35 T, will be necessary. These future technological advancements, along with innovative strategies for incorporating ^19^F labels into proteins, will enable the exploration of protein structures and interactions in cellular environments.

## METHODS

## Materials

4-fluoroindole was purchased from Sigma-Aldrich (Catalog number 457396). Magnesium chloride was purchased from MP Biomedicals (catalog number 209844). All other chemicals were purchased from Sigma-Aldrich and Fisher Scientific. ^15^N-ammonium chloride (^15^NH_4_Cl) and ^13^C_6_-glucose (U-13C6-glucose) were purchased from Cambridge Isotope Laboratories. Human cell culture supplies were purchased from Sigma-Aldrich: media RPMI-1640 (catalog number R8758), fetal bovine serum (catalog number F2442), and A2780 human ovarian adenocarcinoma cell line, and from Corning: Trypsin/EDTA (catalog number 25-052-Cl), Dulbecco’s Phosphate Buffered Saline (catalog number 21-031-CV).

### Sample preparation

#### Protein expression and purification

4F-Trp-U-^13^C,^15^N-CypA was expressed using a protocol similar to that previously described ^26^. Briefly, the CypA gene was inserted into a pET21 vector and *E. coli* Rosetta (DE3) cells were transformed with this vector. The plasmid DNA was sequenced to confirm the amino acid sequence. For protein production, 1 L of modified M9 minimal medium supplemented with 1g/L of ^15^NH_4_Cl and 2 g/L of U-^13^C_6_-glucose was inoculated with ∼25 mls of an overnight culture grown in LuriaBertani Broth to an Absorbance at 600 nm (A600) of 0.1. The cells were grown at 37 °C and at A600 of 0.8, the temperature was reduced to 18 °C, and 25 mg/L of 4F-indole was added to the culture medium. After ∼30-45 minutes, protein expression was induced by adding 0.5 mM IPTG, followed by growth over 16-18 hours. Cells were harvested by centrifugation at 4,000 x g for 5 min at 4 ºC and resuspended in 25 ml lysis buffer (25 mM sodium phosphate, pH 7.0). Cells were lysed by sonication (15 seconds on, 45 seconds off for 5-10 minutes) on ice. Cell debris was removed by centrifugation at 5,000 x g at 4 °C for 15 minutes. The supernatant was collected and further clarified by centrifugation at 17,000 x g at 4 °C for 1 hour. The pH of the clarified lysate was adjusted to pH 5.5 using a 0.1 M acetic acid solution, and any precipitate was removed by centrifugation at 17,000 x g at 4 °C for 1 hour. The supernatant was loaded onto a cation exchange column (SP Cytiva 5 mL column), and protein was eluted using a linear NaCl gradient from 0 to 1 M (Buffer A: 25 mM sodium phosphate, pH 5.5; Buffer B: 25 mM sodium phosphate, 1 M NaCl, pH 5.5). The fractions containing CypA were pooled and further purified by size exclusion chromatography (16/200 Superdex) in 25 mM sodium phosphate buffer, 1 mM DTT, pH 5.5.

The protein quality and extent of labeling with ^15^N, ^13^C, and ^19^F were assessed by mass spectrometry(Figure S1C) and solution NMR ^1^H-^15^N HSQC spectroscopy (Figure S5).

#### A2780 cell culture and protein electroporation

4F-Trp,U-^13^C,^15^N-CypA was delivered into human A2780 cells via electroporation using the procedure developed by the Selenko group ^27^. Before delivery, the protein was concentrated to 2-5 mM, and the buffer was exchanged overnight to 100 mM sodium phosphate, 5 mM KCl, 15 mM MgCl_2_, 15 mM HEPES, 2 mM glutathione reductase, pH 7 (referred to as electroporation buffer, EP). A2780 cells were grown in a T175 flask using RPMI-1640 medium supplemented with 10% fetal bovine serum (FBS) to ∼80% confluency. Cells were detached by trypsin/EDTA treatment (5 mL/plate) for 5 min at 37 ºC. The trypsin was deactivated by adding 25 mL of RPMI medium supplemented with 10% FBS, and cells were collected by centrifugation (150 x g at room temperature for 5 min). The cell pellet was washed twice with 10 mL of pre-warmed Dulbecco’s Phosphate Buffered Saline (DPBS) and once with 1 mL of EP buffer. 1 mL of protein solution in EP buffer was supplemented with 2 mM ATP, passed through a 0.22 μm syringe filter, and added to the A2780 cell pellet. After ∼1 min of incubation, ∼15-20 million cells were transferred to Lonza cuvettes (∼110 μL cell solution/cuvette) and twice electroporated using the B-028 program in the Amaxa Nucleofactor IIb (Lonza Inc.). After electroporation, 1 mL of warm RPMI-1640 medium supplemented with 10% FBS was immediately added to the cuvette, and the content was transferred into a T175 flask containing warm RPMI-1640 medium supplemented with 10% FBS. Cells were placed into the incubator for recovery for 4 to 6 hours. After recovery, dead cells were removed by washing with DPBS three times.. The remaining attached cells were collected using trypsin treatment. Cell viability was assessed using Trypan blue exclusion dye (Sigma-Aldrich) with a Neubauer hemocytometer.

#### In-cell DNP sample preparation

Electroporated cells containing 4F-Trp,U-^13^C,^15^N-CypA were resuspended in RPMI-1640 medium, supplemented with 10% FBS and kept on ice. About 20 million cells were placed into a 1.5 mL Eppendorf and centrifuged at 150 x g for 5 min at 4 °C. The cell pellet was resuspended into a 25 μL solution of 50% RPMI-1640, 20% FBS, 20% D_2_O, 10% DMSO-d_6_, and 30 mM AMUPol (in-cell DNP buffer). The cells were washed twice with 25 μL of the in-cell DNP buffer. Prior to packing the cell into the 3.2 mm DNP MAS NMR rotor, 25 μL of in-cell DNP buffer was added into the DNP rotor. Cells were packed into the rotor by centrifugation at 200 x g for 5 min at 4 °C. Immediately after packing, the rotor was transferred into a homemade Styrofoam container and placed at -80 °C for 24 hours. The cooling container had previously been tested to ensure cell preservation. The final concentration of AMUPol in the sample packed into 3.2 mm rotor was 6.4 mM, as determined by EPR. The measurements were performed on an ESR5000 instrument, at the liquid nitrogen’s boiling point.

#### Protein quantification

After the recovery period after EP, about 1 million of electroporated cells were collected and centrifuged at 300 x g for 5 min. The supernatant was discarded, and cells were resuspended in 30 μL of water. The cells were lysed by 11 freeze-thaw cycles using liquid nitrogen and, subsequently, three cycles of sonication (Branson Sonifier cell disruptor 185) for 3 sec with the output control set at 3 to ensure fragmentation of DNA and RNA. Afterward, 15 μL of the cell lysate was diluted 1 to 1 (volumetric ratio) with SDS-PAGE loading dye, and 5 μL of the mixed sample was loaded onto a 12% SDS-PAGE gel. For protein quantification, we used a calibration curve obtained by running the purified protein at various concentrations (20 to 1 μM) on a SDS-PAGE gel. We estimated the number of moles of protein in the DNP MAS NMR rotor as follows:

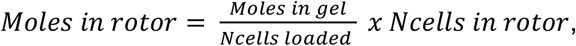

where Ncells loaded are the number of cells loaded onto the gel, and Ncells in the rotor are the number of cells packed into a 3.2 mm DNP MAS NMR rotor (∼20 million cells). The active rotor volume was 30 μL.

### NMR spectroscopy

#### DNP-enhanced MAS NMR experiments

All DNP MAS NMR experiments were performed at 9.4 T (^1^H Larmor frequency of 400.56 MHz) on a Bruker Advance NEO DNP spectrometer equipped with a klystron microwave source operating at 263 GHz electron frequency and with an output power <5 W and a 3.2 mm HFX DNP MAS probe. The MAS frequency was 12 kHz. ^19^F and ^13^C chemical shifts were referenced to mefloquine (with the most downfield peak set at 8.9 ppm) and adamantane, respectively. The in-cell ^1^H DNP buildup time for the 4F-Trp-U-^13^C,^15^N-CypA sample containing 6.4 mM AMUPol was assessed using a pseudo-2D pulse sequence with the following delays: 0.001, 0.01, 0.05, 0.10, 0.50, 1.00, 2.00, 3.00, 5.00, 7.00, 10.00, 15.00, 20.00, 30.00, 45.00, and 60.00 s. For extracting CSA and integrals, all the ^19^F detected spectra were processed using an exponential apodization with 400 Hz line broadening and 400 points, corresponding to an acquisition time of 1.8 ms. The reported SNR^I^, SNR^int^, and SNR^int^_norm_ values were determined from ^1^H-^19^F DNP CPMAS spectra acquired for 4.9 ms and processed without apodization. The (^1^H-^19^F)-^13^C spectrum was acquired for 5.8 ms and processed using an exponential apodization with 100 Hz line broadening. Table S2 and Figure 3 legend summarize other data acquisition and processing parameters.

## ASSOCIATED CONTENT

## Supporting Information

The Supporting Information contains in-cell ^19^F DNP MAS NMR spectra, ^1^H DNP signal enhancements, a solution ^1^H-^15^N HSQC spectrum, ^19^F CSA tensor parameters, acquisition parameters for in-cell DNP MAS experiments, ^13^C chemical shifts of the CypA residues within 6 Å from W121-Hε3, average distances from W121-Hε3 to carbon atoms within 6 Å, and ^13^C chemical shifts of resonances observed in the incell (^1^H-^19^F)-^13^C DNP CPMAS NMR spectrum with tentative residue-specific assignments.

## AUTHOR INFORMATION

### Author Contributions

The manuscript was written through the contributions of all authors. All authors have given approval to the final version of the manuscript. Specific author contributions are given in the Supporting Information.

### Funding Sources

This work was supported by the National Science Foundation (NSF grant MCB2231099 to T.P. and A.M.G.) and the National Institutes of Health (NIH Grant 1U54AI170791 to A.M.G. and T.P., NMR Core).

## ACKNOWLEDGMENT

We thank Armin Purea for supervising the 3.2 mm HFX DNP probe production.

## ABBREVIATIONS

### NOMENCLATURE

NMR: nuclear magnetic resonance
MAS: magic-angle spinning
DNP: dynamic nuclear polarization
CypA: cyclophilin A
CP: cross polarization

## Notes

### Competing Interest Statement

The authors have declared no competing interest.

## REFERENCES

(1) Beck, M.; Covino, R.; Hänelt, I.; Müller-McNicoll, M. Understanding the Cell: Future Views of Structural Biology. Cell 2024, 187 (3), 545–562. 10.1016/j.cell.2023.12.017.

(2) Serber, Z.; Keatinge-Clay, A. T.; Ledwidge, R.; Kelly, A. E.; Miller, S. M.; Dötsch, V. High-Resolution Macromolecular NMR Spectroscopy inside Living Cells. J. Am. Chem. Soc. 2001, 123 (10), 2446–2447. 10.1021/ja0057528.

(3) Selenko, P.; Serber, Z.; Gadea, B.; Ruderman, J.; Wagner, G. Quantitative NMR Analysis of the Protein GB1 Domain in Xenopus Laevis Egg Extracts and Intact Oocytes. Proc. Natl. Acad. Sci. 2006, 103 (32), 11904–11909. 10.1073/pnas.0604667103.

(4) Inomata, K.; Ohno, A.; Tochio, H.; Isogai, S.; Tenno, T.; Nakase, I.; Takeuchi, T.; Futaki, S.; Ito, Y.; Hiroaki, H.; Shirakawa, M. High-Resolution Multi-Dimensional NMR Spectroscopy of Proteins in Human Cells. Nature 2009, 458 (7234), 106–109. 10.1038/nature07839.

(5) Zhu, W.; Guseman, A. J.; Bhinderwala, F.; Lu, M.; Su, X.; Gronenborn, A. M. Visualizing Proteins in Mammalian Cells by 19F NMR Spectroscopy. Angew. Chem. Int. Ed. 2022, 61 (23), e202201097. 10.1002/anie.202201097.

(6) Narasimhan, S.; Scherpe, S.; Lucini Paioni, A.; Zwan, J.; Folkers, G. E.; Ovaa, H.; Baldus, M. DNP-supported Solid-state NMR Spectroscopy of Proteins inside Mammalian Cells. Angew. Chem. Int. Ed. 2019, 58 (37), 12969–12973. 10.1002/anie.201903246.

(7) Movellan, K. T.; Zhu, W.; Banks, D.; Kempf, J.; Runge, B.; Gronenborn, A. M.; Polenova, T. Expanding the Tool Box for Native Structural Biology:19F Dynamic Nuclear Polarization with Fast Magic Angle Spinning. Sci. Adv. 2024, 10 (40), eadq3115. 10.1126/sciadv.adq3115.

(8) Costello, W. N.; Xiao, Y.; Mentink-Vigier, F.; Kragelj, J.; Frederick, K. K. DNP-Assisted Solid-State NMR Enables Detection of Proteins at Nanomolar Concentrations in Fully Protonated Cellular Milieu. J. Biomol. NMR 2024, 78 (2), 95–108. 10.1007/s10858-024-00436-9.

(9) Biedenbänder, T.; Aladin, V.; Saeidpour, S.; Corzilius, B. Dynamic Nuclear Polarization for Sensitivity Enhancement in Biomolecular Solid-State NMR. Chem. Rev. 2022, 122 (10), 9738–9794. 10.1021/acs.chemrev.1c00776.

(10) Theillet, F.-X. In-Cell Structural Biology by NMR: The Benefits of the Atomic Scale. Chem. Rev. 2022, 122 (10), 9497–9570. 10.1021/acs.chemrev.1c00937.

(11) Overall, S. A.; Barnes, A. B. Biomolecular Perturbations in In-Cell Dynamic Nuclear Polarization Experiments. Front. Mol. Biosci. 2021, 8, 743829. 10.3389/fmolb.2021.743829.

(12) Ghosh, R.; Xiao, Y.; Kragelj, J.; Frederick, K. K. In-Cell Sensitivity-Enhanced NMR of Intact Viable Mammalian Cells. J. Am. Chem. Soc. 2021, 143 (44), 18454–18466. 10.1021/jacs.1c06680.

(13) Beriashvili, D.; Yao, R.; D’Amico, F.; Krafčíková, M.; Gurinov, A.; Safeer, A.; Cai, X.; Mulder, M. P. C.; Liu, Y.; Folkers, G. E.; Baldus, M. A High-Field Cellular DNP-Supported Solid-State NMR Approach to Study Proteins with Sub-Cellular Specificity. Chem. Sci. 2023, 14 (36), 9892–9899. 10.1039/D3SC02117C.

(14) Bertarello, A.; Berruyer, P.; Artelsmair, M.; Elmore, C. S.; Heydarkhan-Hagvall, S.; Schade, M.; Chiarparin, E.; Schantz, S.; Emsley, L. In-Cell Quantification of Drugs by Magic-Angle Spinning Dynamic Nuclear Polarization NMR. J. Am. Chem. Soc. 2022, 144 (15), 6734–6741. 10.1021/jacs.1c12442.

(15) Pham, L. B. T.; Costantino, A.; Barbieri, L.; Calderone, V.; Luchinat, E.; Banci, L. Direct Expression of Fluorinated Proteins in Human Cells for 19F in-Cell NMR Spectroscopy. J. Am. Chem. Soc. 2023, 145 (2), 1389–1399. 10.1021/jacs.2c12086.

(16) Šoltésová, M.; Pinon, A. C.; Aussenac, F.; Schlagnitweit, J.; Reiter, C.; Purea, A.; Melzi, R.; Engelke, F.; Martin, D.; Krambeck, S.; Biscans, A.; Kay, E.; Emsley, L.; Schantz, S. 1H–19F Cross-Polarization Magic Angle Spinning Dynamic Nuclear Polarization NMR Investigation of Advanced Pharmaceutical Formulations. J. Magn. Reson. 2025, 371, 107827. 10.1016/j.jmr.2024.107827.

(17) Zhang, X. C.; Xue, K.; Salvi, M.; Schomburg, B.; Mehrens, J.; Giller, K.; Stopp, M.; Weisenburger, S.; Böning, D.; Sandoghdar, V.; Unden, G.; Becker, S.; Andreas, L. B.; Griesinger, C. Mechanism of Sensor Kinase CitA Transmembrane Signaling. Nat. Commun. 2025, 16 (1), 935. 10.1038/s41467-024-55671-3.

(18) Gerig, J. Fluorine NMR of Proteins. Prog. Nucl. Magn. Reson. Spectrosc. 1994, 26, 293–370.

(19) Gronenborn, A. M. Small, but Powerful and Attractive: 19F in Biomolecular NMR. Structure 2022, 30 (1), 6–14. 10.1016/j.str.2021.09.009.

(20) Quinn, C. M.; Zadorozhnyi, R.; Struppe, J.; Sergeyev, I. V.; Gronenborn, A. M.; Polenova, T. Fast 19F Magic-Angle Spinning Nuclear Magnetic Resonance for the Structural Characterization of Active Pharmaceutical Ingredients in Blockbuster Drugs. Anal. Chem. 2021, 93 (38), 13029–13037. 10.1021/acs.analchem.1c02917.

(21) Runge, B. R.; Zadorozhnyi, R.; Quinn, C. M.; Russell, R. W.; Lu, M.; Antolínez, S.; Struppe, J.; Schwieters, C. D.; Byeon, I.-J. L.; Hadden-Perilla, J. A.; Gronenborn, A. M.; Polenova, T. Integrating 19F Distance Restraints for Accurate Protein Structure Determination by Magic Angle Spinning NMR Spectroscopy. J. Am. Chem. Soc. 2024, 146 (44), 30483–30494. 10.1021/jacs.4c11373.

(22) Kalabekova, R.; Quinn, C. M.; Movellan, K. T.; Gronenborn, A. M.; Akke, M.; Polenova, T. 19F Fast Magic-Angle Spinning NMR Spectroscopy on Microcrystalline Complexes of Fluorinated Ligands and the Carbohydrate Recognit Ion Domain of Galectin-3. Biochemistry 2024, 63 (17), 2207–2216. 10.1021/acs.biochem.4c00232.

(23) Lu, M.; Wang, M.; Sergeyev, I. V.; Quinn, C. M.; Struppe, J.; Rosay, M.; Maas, W.; Gronenborn, A. M.; Polenova, T. 19F Dynamic Nuclear Polarization at Fast Magic Angle Spinning for NMR of HIV-1 Capsid Protein Assemblies. J. Am. Chem. Soc. 2019, 141 (14), 5681–5691. 10.1021/jacs.8b09216.

(24) Lu, M.; Sarkar, S.; Wang, M.; Kraus, J.; Fritz, M.; Quinn, C. M.; Bai, S.; Holmes, S. T.; Dybowski, C.; Yap, G. P. A.; Struppe, J.; Sergeyev, I. V.; Maas, W.; Gronenborn, A. M.; Polenova, T. 19F Magic Angle Spinning NMR Spectroscopy and Density Functional Theory Calculations of Fluorosubstituted Tryptophans: Integrating Experiment and Theory for Accurate Determination of Chemical Shift Tensors. J. Phys. Chem. B 2018, 122 (23), 6148–6155. 10.1021/acs.jpcb.8b00377.

(25) Albert, B. J.; Gao, C.; Sesti, E. L.; Saliba, E. P.; Alaniva, N.; Scott, F. J.; Sigurdsson, S. Th.; Barnes, A. B. Dynamic Nuclear Polarization Nuclear Magnetic Resonance in Human Cells Using Fluorescent Polarizing Agents. Biochemistry 2018, 57 (31), 4741–4746. 10.1021/acs.biochem.8b00257.

(26) Lu, M.; Ishima, R.; Polenova, T.; Gronenborn, A. M. 19F NMR Relaxation Studies of Fluorosubstituted Tryptophans. J. Biomol. NMR 2019, 73 (8–9), 401–409. 10.1007/S10858-019-00268-Y.

(27) Theillet, F.-X.; Binolfi, A.; Bekei, B.; Martorana, A.; Rose, H. M.; Stuiver, M.; Verzini, S.; Lorenz, D.; Van Rossum, M.; Goldfarb, D.; Selenko, P. Structural Disorder of Monomeric α-Synuclein Persists in Mammalian Cells. Nature 2016, 530 (7588), 45–50. 10.1038/nature16531.

