## Supporting Information for "Pushing Sensitivity and Specificity Limits in Native Structural Biology: ^19^F Multinuclear Dynamic Nuclear Polarization with Magic Angle Spinning"

### **Table S1.** ^19^F CSA tensor parameters for 4F-Trp,U-^13^C,^15^N-CypA delivered into human A2780 cells.

|  | Integrated intensity, relative to isotropic peak (%) | Integrated intensity, normalized |
| --- | --- | --- |
| Isotropic peak | 100 | 0.48 |
| 1^st^ ssb (shielded) | 30 | 0.14 |
| 2^nd^ ssb (shielded) | 15 | 0.07 |
| 3^rd^ ssb (shielded) | 2 | 0.01 |
| -1^st^ ssb (deshielded) | 50 | 0.24 |
| -2^nd^ ssb (deshielded) | 10 | 0.05 |
| -3^rd^ ssb (deshielded) | 1 | 0.01 |
| δ_σ_ (ppm) | 67±4 | |
| η | 0.5±0.4 | |

### **Table S2.** Acquisition parameters for in-cell DNP MAS NMR experiments on 4F-Trp,U-^13^C,^15^N-CypA.

| **Experiment** | **^19^F direct excitation** | **^1^H-^19^F**  **CPMAS** | **^1^H-^13^C**  **CPMAS** | **(^1^H-^19^F)-^13^C**  **CPMAS** |
| --- | --- | --- | --- | --- |
| **RF field (kHz)** |  |  |  |  |
| ^1^H |  | 71 | 71 | 71 |
| ^19^F | 71 | 71 |  | 71 |
| ^13^C |  |  | 56 | 56 |
| **^1^H decoupling RF field (kHz)** |  |  |  |  |
| ^1^H | 55 | 55 | 55 | 55 |
| **1^st^ CP transfer (^1^H to ^13^C/^19^F)** |  |  |  |  |
| ^1^H (kHz) |  | 55-46 | 55-46 | 55-46 |
| ^13^C/^19^F (kHz) |  | 64 |  | 59 |
| Contact time (ms) |  | 0.5 |  | 0.5 |
| **2^nd^ CP transfer (^19^F to ^13^C)** |  |  |  |  |
| ^19^F (kHz) |  |  |  | 90 |
| ^13^C (kHz) |  |  |  | 43-35.77 |
| Contact time (ms) |  |  |  | 0.5 |
| **Spectral width (kHz)** |  |  |  |  |
| ^19^F | 114 | 114 |  |  |
| ^13^C |  |  | 114 | 96 |
| **Acquisition time** |  |  |  |  |
| t_1_ (ms) / points | 4.88 / 1110 | 4.88 / 1110 | 4.88 / 1110 | 5.77 (1110) |
| **Number of scans** |  |  |  |  |
| μw ON | 4096 | 2048 | 256 | 61440 |
| μw OFF |  | 8192 | 4096 |  |
| **Experimental time (min)** |  |  |  |  |
| μw ON | 276 | 103 | 13 | 3072 |
| μw OFF |  | 411 | 205 |  |

### **Table S3.** ^13^C chemical shifts of CypA residues located within 6 Å from around 4F-Trp CypA (BMRB entry BMR27265).

| **Residue** | **Chemical shift (ppm)** | | | | | | |
| --- | --- | --- | --- | --- | --- | --- | --- |
|  | **Co** | **Cα** | **Cβ** | **Cγ** | **Cδ (δ1/δ2)** | **Cε (ε2/ε3)** | **Cζ (ζ2/ζ3)** |
| **K118** | 177.2 |  |  | 25 |  |  |  |
| **T119** | 177.9 | 57.2 | 68.7 | 19.9 |  |  |  |
| **E120** | 178.2 | 59.4 | 29.2 | 35.5 | - |  |  |
| **W121** | 177.2 | 59.8 | 26.95 |  |  |  |  |

### **Table S4.** Average distances from W121-Hε3 (position 4) to carbon atoms within 6 Å from the X-ray structure of CypA (PDB: 3K0M).

|  | **Co** | **Cα** | **Cβ** | **Cγ** | **Cδ (δ1)** |
| --- | --- | --- | --- | --- | --- |
| **K118** | 5.5 |  |  |  |  |
| **T119** | 4.0 | 4.3 | 5.2 | 5.9 |  |
| **E120** | 4.1 | 4.3 | 4.7 | 3.9 | 3.3 |
| **W121** | 4.9 | 3.7 | 3.1 |  |  |

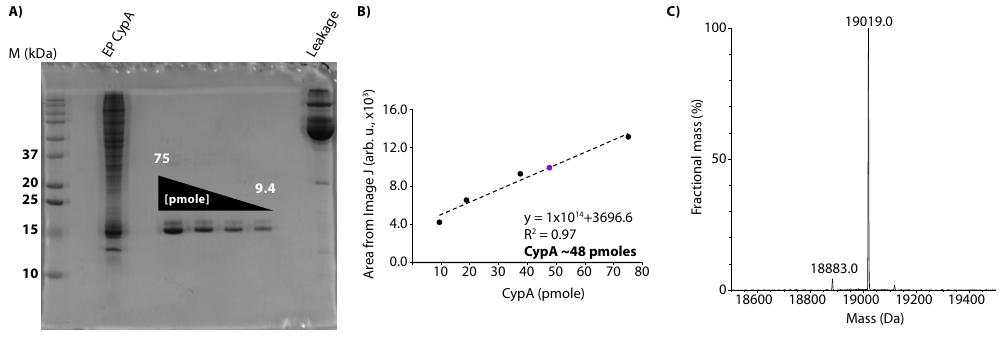

### **Figure S1. Protein quantification and fluorine incorporation levels in CypA**. **(A)** SDS-PAGE shows successful delivery and the absence of protein leakage while packing the MAS NMR rotor for in-cell CypA sample. **(B)** Protein quantification was obtained by extracting the intensities from SDS-PAGE using ImageJ. **(C)** The mass spectrum shows a single species with a mass of 19019 Da (theoretically expected mass is 19041 Da), indicating a fluorine incorporation level above 90%.

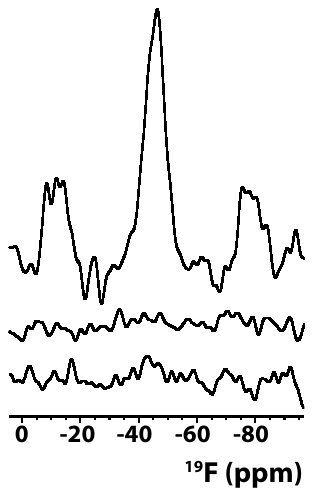

### **Figure S2. In-cell ^19^F-detected DNP MAS NMR spectra of 4F-Trp,U-^13^C,^15^N-CypA.** ^1^H-^19^F DNP-enhanced CPMAS spectra with microwave on (top) and off (middle). ^19^F direct-excitation DNP MAS NMR spectrum with microwave on (bottom). The CPMAS spectra were recorded with 2048 scans for a total of 103 min and 8192 scans for a total of 411 min for microwave on and off, respectively. The direct-excitation spectrum was recorded with 4096 scans for a total of 276 min.

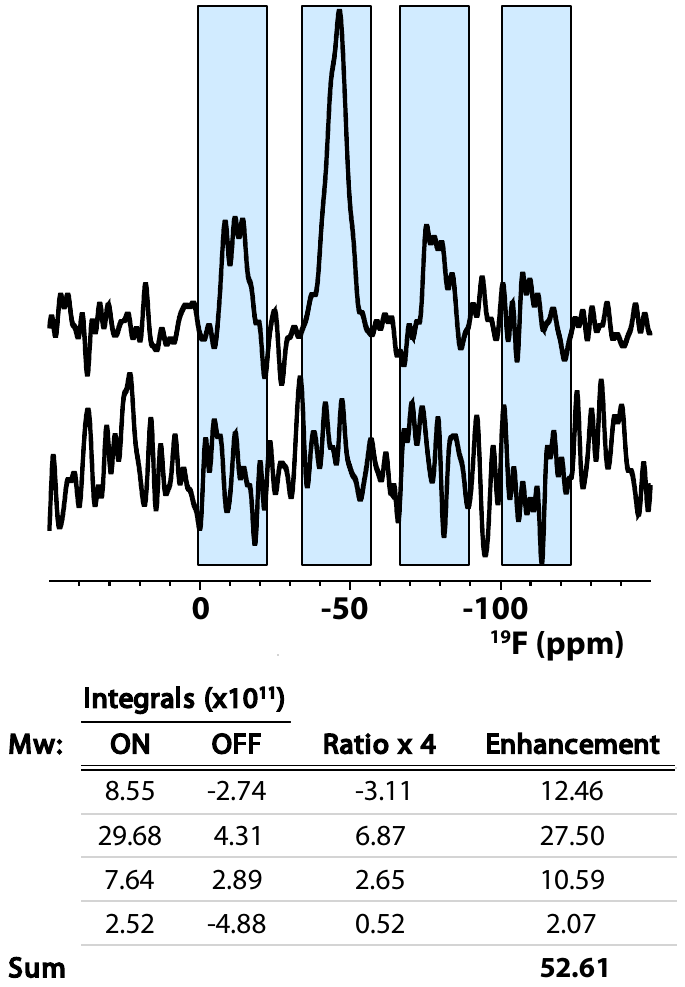

### **Figure S3. ^1^H-^19^F in-cell DNP CPMAS signal enhancements.** ^1^H-^19^F CPMAS spectra acquired with microwaves on (top) and off (bottom). To estimate DNP enhancements, the ratio of the integrated signal intensities of both spectra was calculated using spectral regions containing signal in the DNP-enhanced spectrum (blue). The spectra were recorded with 2048 scans for a total of 103 min (microwaves on) and 8192 scans for a total of 411 min (microwaves off).

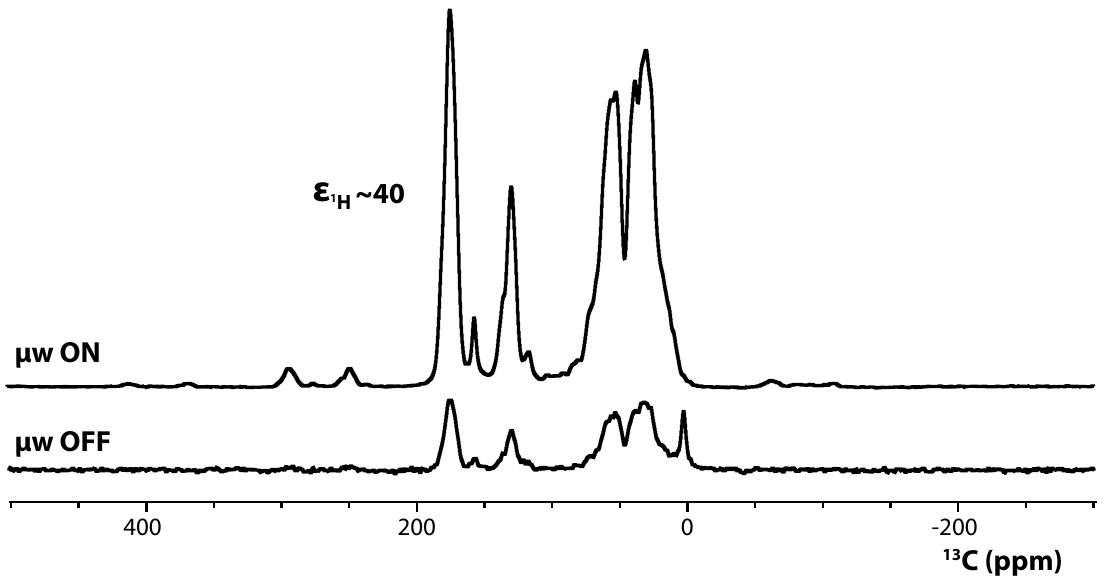

### **Figure S4. DNP signal enhancements in in-cell ^1^H-^13^C CPMAS experiments.** In-cell ^1^H-^13^C DNP-enhanced CPMAS NMR spectra of 4F-Trp,U-^13^C,^15^N-CypA with microwave off (bottom, 4,096 scans, ~3.5 hours) and on (top, 512 scans, 13 min).

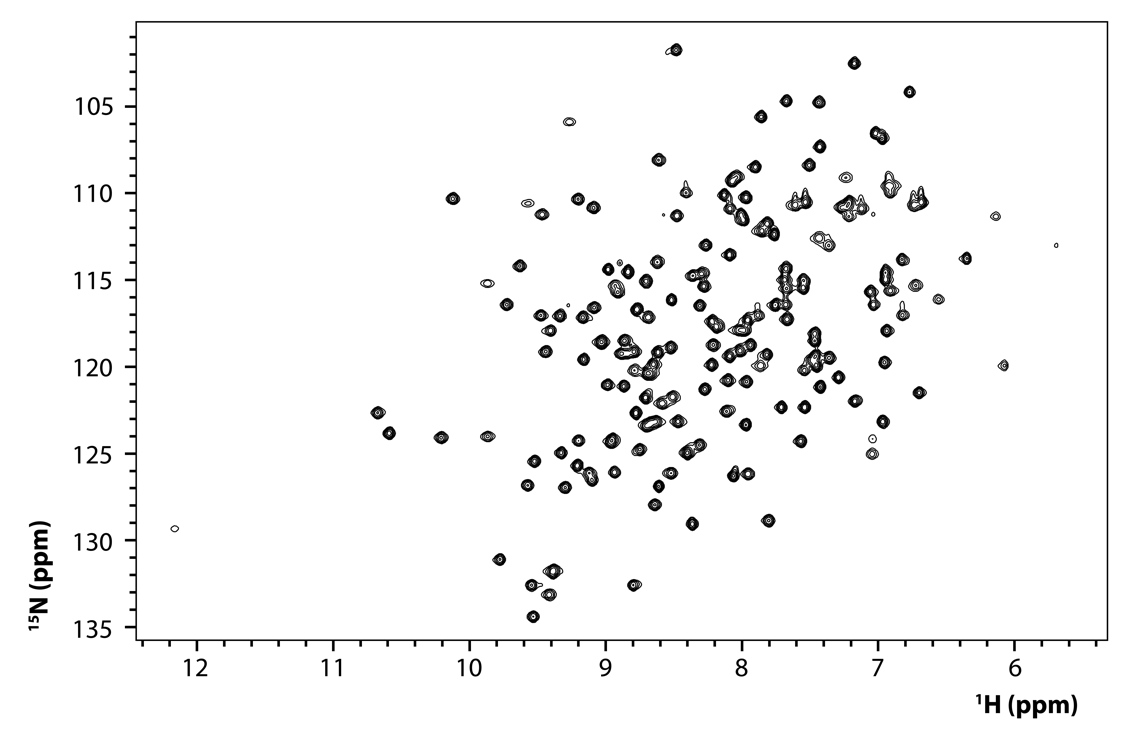

### **Figure S5. ^1^H-^15^N-HSQC spectrum of 4F-Trp,U-^13^C,^15^N-CypA in EP buffer.** The spectrum demonstrates that the ^19^F incorporation at position 4 of the Trp side chain does not affect the protein fold. The NMR ^15^NεH signal (^1^Hε1=9.6 ppm / ^15^Nε1=130.1 ppm) from the Trp-121 side chain is essentially undetectable, indicating that ^19^F incorporation level is above 90%.

### **AUTHOR CONTRIBUTIONS**

**Conceptualization**: TP and AMG; **Methodology**: TP, AMG, KTM; **Investigation**: KTM, TP, DB; **Data interpretation**: KTM and TP; **Supervision**: TP, AMG, JK; **Resources**: TP, DB, KTM, JK, CR; **Writing**: *original draft*: TP and KTM, *review & editing*: TP, AMG, KTM, DB, JK; **Project administration**: TP, AMG, JK; **Funding acquisition**: TP and AMG
